# Crossmodal Hierarchical Predictive Coding for Audiovisual Sequences in Human Brain

**DOI:** 10.1101/2023.11.19.567767

**Authors:** Yiyuan Teresa Huang, Chien-Te Wu, Yi-Xin Miranda Fang, Chin-Kun Fu, Shinsuke Koike, Zenas C. Chao

## Abstract

Predictive-coding theory proposes that the brain actively predicts sensory inputs based on prior knowledge. While this theory has been extensively researched within individual sensory modalities, there is a crucial need for empirical evidence supporting hierarchical predictive processing across different modalities to further generalize the theory. Here, we examine how crossmodal knowledge is represented and learned in the brain by identifying the hierarchical networks underlying crossmodal predictions when information of one sensory modality leads to a prediction in another modality. We record electroencephalogram (EEG) in humans during a crossmodal audiovisual local-global oddball paradigm, in which the predictability of transitions between tones and images are manipulated at two hierarchical levels: stimulus-to-stimulus transition (local level) and multi-stimulus sequence structure (global level). With a model-fitting approach, we decompose the EEG data using three distinct predictive-coding models: one with no audiovisual integration, one with audiovisual integration at the global level, and one with audiovisual integration at both the local and global levels. The best-fitting model demonstrates that audiovisual integration occurs at both levels. This highlights a convergence of auditory and visual information to construct crossmodal predictions, even in the more basic interactions that occur between individual stimuli. Furthermore, we reveal the spatio-spectro-temporal signatures of prediction-error signals across hierarchies and modalities, and show that auditory and visual prediction-error signals are progressively redirected to the central-parietal area of the brain as learning progresses. Our findings unveil a crossmodal predictive coding mechanism, where the unimodal framework is implemented through more distributed brain networks to process hierarchical crossmodal knowledge.

## Introduction

Predictive-coding theory postulates that the brain is constantly predicting upcoming events based on an internal model established by learning the statistical regularity in sensory experience (Clark, 2013; Friston et al., 2012). This is achieved by a bidirectional and hierarchical signaling cascade, where the top-down prediction represents the information inferred from prior knowledge to explain away the input, and the bottom-up prediction error is generated to refine the prediction when there is an unexplained discrepancy. This coding is thought to be the basis for diverse cognitive processes, such as perceptual inference, reinforcement learning, and motivational attention (den Ouden et al., 2012; Friston, 2018; Knill & Pouget, 2004).

Considerable evidence of predictive coding has been provided within single sensory modalities. For example, the mismatch negativity (MMN), a surprise response to oddball stimuli, has been interpreted as a prediction-error signal in the domains of audition (Näätänen, 1995; Näätänen et al., 2007; Wacongne et al., 2012), vision (Heslenfeld, 2003; Stefanics et al., 2014) and tactility (Shen et al., 2018; Shinozaki et al., 1998). The hierarchical organization of the unimodal predictive coding has also been examined, for example, by the local-global oddball paradigm with auditory (Bekinschtein et al., 2009) and visual stimuli (Blundon et al., 2017). In this paradigm, regularities were controlled at the stimulus-to-stimulus level (local regularity) and the multi-stimulus sequence level (global regularity). The hierarchical prediction-error signals have been identified in both monkeys and humans (Chao et al., 2018; El Karoui et al., 2015; Wacongne et al., 2011), and hierarchical prediction signals have been extracted with a quantitative predictive-coding model (Chao et al., 2022).

To further generalize the predictive-coding theory, it is critical to verify how predictive coding can operate across multiple sensory modalities, particularly, how prior knowledge can be transferred across sensory domains. During audiovisual events, it has been found that visual information can facilitate the early auditory response, suggesting a prediction established by the crossmodal transition from the visual to auditory domains and thus speeding up the process of upcoming auditory input (van Wassenhove et al., 2005; Vroomen & Stekelenburg, 2010).

Furthermore, the concept of a crossmodal prediction error comes into play once a crossmodal transition is violated. For instance, a visual stimulus V followed by an auditory stimulus A is learned, and a crossmodal prediction error is generated when this expected transition is disrupted by replacing A with a different auditory stimulus A’. This error, which arises when visual information is inadequate to infer upcoming auditory stimuli, serves to explain phenomena such as the McGurk fusion and combination of audiovisual speech (Olasagasti et al., 2015; van Wassenhove et al., 2005). However, it remains unclear how crossmodal transitions are predicted at different hierarchical levels, and how sensory information from different modalities converges along the hierarchy that leads to the emergence of crossmodal prediction errors.

To address this, we use a crossmodal audiovisual local-global paradigm in which auditory and visual stimuli are delivered one at a time in sequences, and the unimodal and crossmodal transitions are controlled at two hierarchical levels of temporal regularities. By delivering stimuli one at a time, we can independently control the occurrences of both unimodal and crossmodal transitions, which cannot be done if both auditory and visual stimuli are presented simultaneously. To examine how the brain learns and represents unimodal and crossmodal transitions at the two levels, we record electroencephalogram (EEG) and analyze the data with a model-fitting approach (Chao et al., 2022). Three predictive-coding models are tested: one with no audiovisual integration, one with audiovisual integration at the global level, and one with audiovisual integration at both the local and global levels. The best-fitting model reveals that audiovisual integration occurs at both levels, indicating a convergence of the auditory and visual information to create crossmodal predictions, even at the lower level of stimulus-to-stimulus transitions. We further show that the auditory and visual prediction-error signals are gradually re-routed from unimodal areas to the central-parietal area during learning. Our findings demonstrate that the principles of hierarchical predictive coding, initially understood in the context of unimodal processing, can also comprehensively describe the learning and encoding of multilevel statistics observed in crossmodal transitions, with the engagement of more widespread brain networks.

## Results

### A Crossmodal Audiovisual Local-Global Oddball Paradigm

We designed a crossmodal audiovisual local-global oddball paradigm in which crossmodal transitions were created between a tone (denoted as A) and a checkerboard (denoted as V). These two stimuli were arranged to create 4 sequences: AAA, AAV, VVV, and VVA. Within each sequence, all stimuli were presented for 200 ms with a 300 ms stimulus onset asynchrony. A random interval of 1600-1900 ms was inserted between the offset of one sequence and the onset of the next sequence (Figure 1A). Four sequence blocks were created, each one with 120 trials of stimulation sequences but with distinct predictability in the crossmodal transitions. In each block, a frequent sequence (100 trials) and an infrequent sequence (20 trials) were assigned, with the first two stimuli in both sequences being of the same modality and the last being of different modalities (Figure 1B). Each block was presented twice: once with both the visual and auditory stimuli delivered from the left side, and once with both stimuli delivered from the right side (Figure 1A). The purpose of this lateral delivery was to enable post-hoc confirmation of the lateralization in sensory responses, such as whether EEG responses to an image in the left visual field would appear in the right occipital area. Participants were asked to pay attention to the stimuli while maintaining eye fixation on a white dot. During the experiment, 64-channel EEG signals were recorded, and no task-related behavioral responses were required.

**Figure 1.**
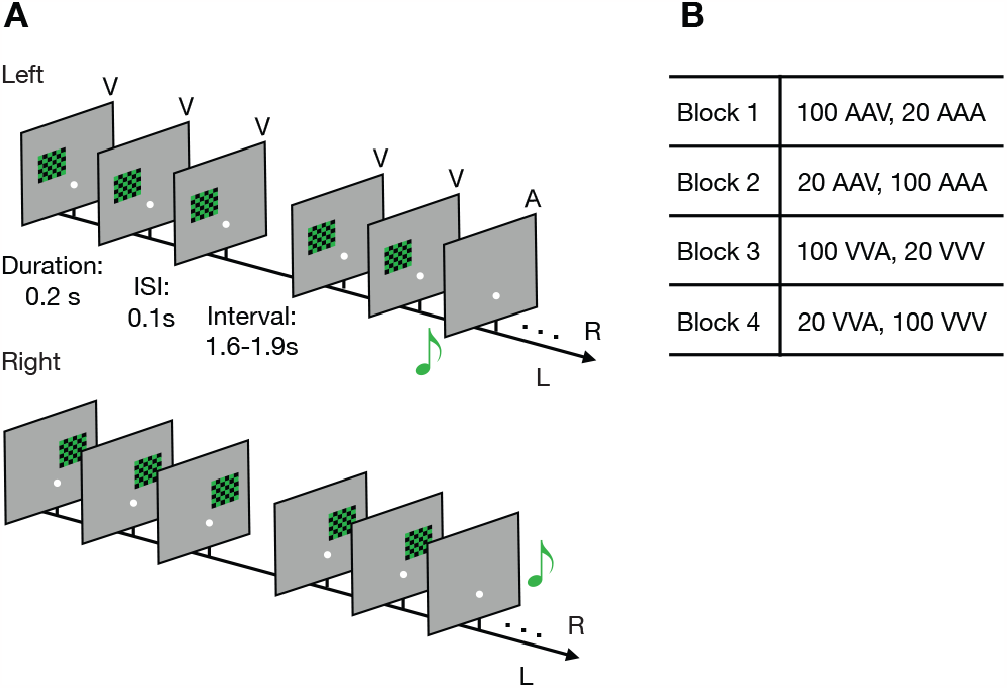
Crossmodal audiovidual local-global oddball paradigm. **(A)** Demonstration of the left-and right-side stimulation delivery. The white dot represents the fixation. The visual stimuli are displayed on left or right relative to the fixation, and the auditory stimuli are presented on the left or right speaker. ISI: inter-stimulus interval. **(B)** Configuration of trial number and trial types in the four blocks. A represents a tone and V represents a green-and-black checkerboard. A and V are used to form three-stimulus sequences.

### Three Hypotheses of Crossmodal Transitions

In the paradigm, the regularities of crossmodal transitions can be established at two hierarchical levels within a single block. The local regularity is established based on the stimulus-to-stimulus transition probability (TP), defined as the conditional probability of a particular stimulus occurring, given the preceding stimulus. We denoted the probabilities of the crossmodal A-to-V and V-to-A transitions as TP_A**→**V_ and TP_V**→**A_, respectively. On the other hand, the probabilities of the unimodal A-to-A and V-to-V transitions were denoted as TP_A**→**A_ and TP_V**→**V_, respectively. At a higher level, the global regularity is established based on the sequence probability (SP), defined as the probability of sequences AAA, AAV, VVV, or VVA within a block. These were denoted as SP_AAA_, SP_AAV_, SP_VVV_, and SP_VVA_, respectively.

To understand how the brain processes local and global regularities in the crossmodal sequences, we first need to know how it integrates auditory and visual information. For example, if there’s no audiovisual integration occurring between two consecutive stimuli (local integration), the probabilities of A-to-V and V-to-A transitions cannot be determined, resulting in TP_A**→**V_ and TP_V**→**A_ being zero. Similarly, without audiovisual integration at the sequence level (global integration), AAV and VVA cannot be recognized as single sequences, and therefore, SP_AAV_ and SP_VVA_ are zero. Here, we present three hypotheses on how the brain integrates auditory and visual information at the local and global levels, which were subsequently verified using EEG data. To elucidate these hypotheses, we use the processing sequences of AAA and AAV as illustrative examples (Figure 2A).

**Figure 2.**
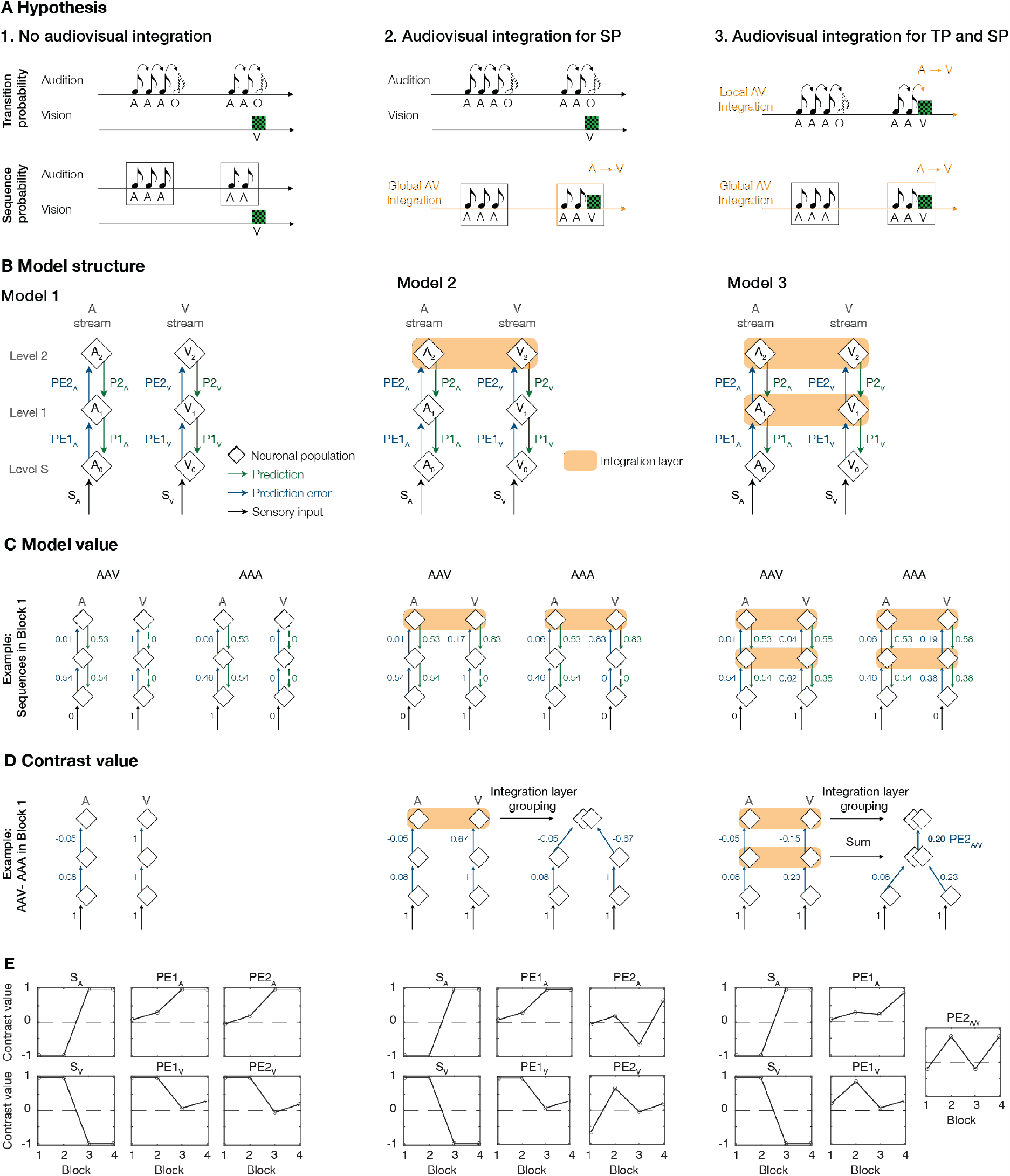
Hypothesis and Crossmodal hierarchical predictive coding models. **(A)** Three hypotheses of how crossmodal transitions are considered in probabilities, shown with the example of AAA and AAV sequences. The transition probability is calculated with the transitions between the stimuli or omission. The black curved arrow represents the transition between the stimuli with the same modality or between a stimulus and omission, the orange curved arrow represents the transition between the stimuli with different modalities. The sequence probability is calculated with the occurrences of the sequence types. The black rectangle rim represents the sequence type composed of an identical modality, while the orange rectangle rim represents the sequence type composed of different modalities. The streams of audition and vision represent the independent processes of auditory and visual stimuli. The stream of audiovisual integration highlighted in the orange solid line represents that stimuli of the two modalities are processed together; that is, the crossmodal transitions can be learned. **(B)** The model structures are built upon the three hypotheses. Each model consists of two sensory streams (A and V) and three levels (S, 1, and 2). For neuronal populations (white diamonds), A_0_, A_1_, A_2_ were denoted as populations in the auditory stream at the level S, level 1, and level 2, respectively. V_0_, V_1_, V_2_ were denoted as populations in the visual stream at the level S, level 1, and level 2, respectively. Prediction signals (green solid downward arrows), prediction error signals (blue solid upward arrows), and sensory inputs (black solid upward arrows) are shown. The horizontal orange bar represents the integration between two streams. **(C)** The model values of the three hypotheses. The example of the model values for AAV and AAA sequences in Block 1 are shown. The values are calculated when the last stimulus of the sequence (underlined) is received. The dashed arrow represents the signal with a zero value. **(D)** The contrast values in Block 1 (first row) are obtained by comparing the model values of AAV to AAA shown in panel A. The green solid lines are removed since no prediction signals are left after the contrast. The two white diamonds being overlaid represent the neuronal populations grouped in the integration layer, and the values are combined by summation when the signals are sent from and to an integration layer. **(E)** The model contrast values of three models in four blocks.

#### Hypothesis 1

Auditory and visual information are not integrated, but instead are processed independently within their respective auditory and visual streams. Since the auditory and visual streams are independent from each other, the auditory stream perceives AAA and AAV as AAA and AA, respectively (Figure 2A). In this scenario, TP_A**→**V_ equals zero, and TP is determined based on the A-to-A transition (TP_A**→**A_) and A-to-O transition (TP_A**→**O_), where O represents omissions at the end of the sequence. It is important to note that the A-to-O transition needs to be considered, since at the stimulus-to-stimulus transition level, the last A in a sequence will continue to predict the next stimulus item (including omission) (Chao et al., 2022). Similarly, SP_AAV_ equals zero, and SP applies only to sequences AAA and AA (thus SP_AAA_ and SP_AA_ are nonzero).

#### Hypothesis 2

Auditory and visual information are not integrated at the local level, but are integrated later at the global level. In this scenario, auditory and visual information are processed separately to determine TP, as in Hypothesis 1, but are processed jointly to determine SP. Consequently, AAA and AAV are considered as sequences (thus SP_AAA_ and SP_AAV_ are nonzero), and there is no sequence AA (SP_AA_ is equal to zero).

#### Hypothesis 3

Auditory and visual information are integrated at both the local and global levels. In this scenario, auditory and visual information are combined to determine both TP and SP. Consequently, the A-to-V transition is now considered, making TP_A**→**V_ nonzero. SP is determined in the same manner as in Hypothesis 2.

Therefore, the TP and SP values vary not only across the blocks but also across the three hypotheses, as detailed in Table 1.

**Table 1.**
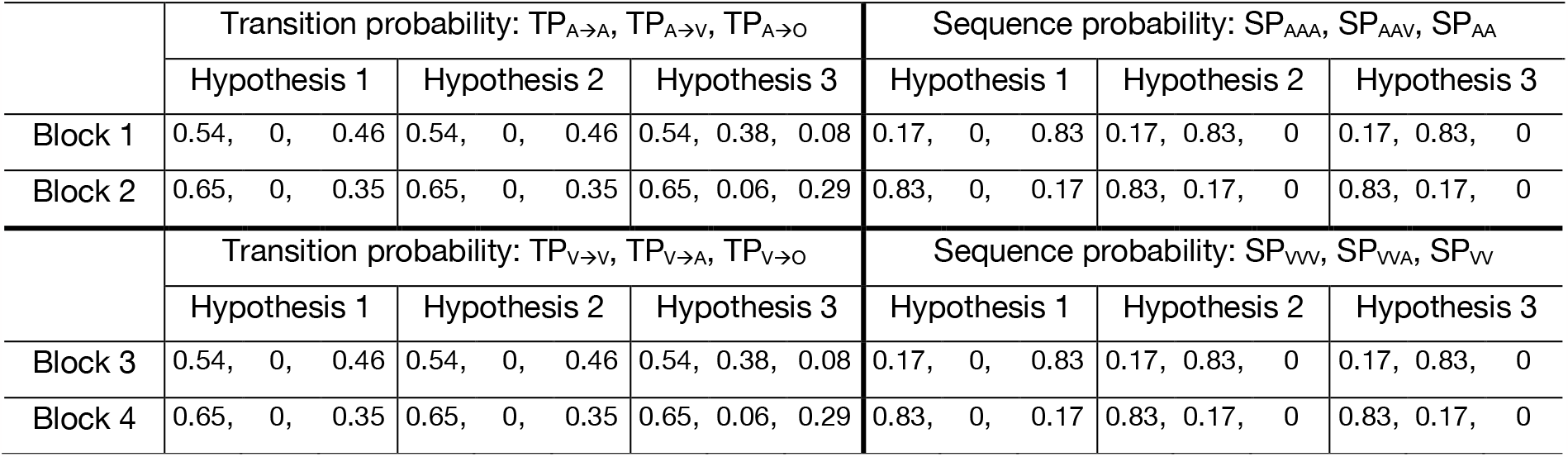
Transition and sequence probabilities across blocks and models.

### Quantitative Models for Evaluating the Hypotheses

In order to evaluate these hypotheses using EEG data, we need to quantify the interplay among sensory, prediction, and prediction-error signals within the varying configurations of audiovisual integration proposed in each hypothesis. To achieve this, we adapted a hierarchical predictive-coding model originally designed to elucidate the auditory local-global paradigm (Chao et al., 2018, 2022), and incorporated varying interactions between auditory and visual streams. Each of the three hypotheses was represented by a corresponding model, each featuring unique structural elements and signal flows (Figure 2B).

All three models consist of two key structures: the auditory and visual sensory streams (A and V streams), and three hierarchical levels (Levels S, 1, and 2). Level S receives sensory input, and Levels 1 and 2 encode the local and global regularities, respectively. Using the A stream as an example, the neuronal population at Level S (denoted as A_0_) receives an auditory input (denoted as S_A_) and a top-down first-level prediction of that input (denoted as P1_A_) from Level 1. When the stimulus is A, S_A_ = 1, and otherwise, S_A_ = 0. The first-level prediction error (denoted as PE1_A_), which is the absolute difference between S_A_ and P1_A_, is sent to Level 1. At Level 1, the neuronal population (denoted as A_1_) receives PE1_A_ and a second-level prediction (denoted as P2_A_) from Level 2. The second-level prediction error (denoted as PE2_A_), which is the absolute difference between PE1_A_ and P2_A_, is sent to the neural population at Level 2 (denoted as A_2_). The V stream functions similarly, and there are first-level and second-level predictions and prediction errors (denoted as P1_V_, P2_V_, PE1_V_, and PE2_V_). S_V_ is set to 1 when the stimulus is V and 0 otherwise.

The key distinction among the three models is whether auditory and visual streams are integrated at Level 1 (local level) and Level 2 (global level) (Figure 2B):

#### Model 1

To model Hypothesis 1, which proposes no audiovisual integration, this model features discrete auditory and visual streams, and the auditory and visual signals operate independently.

#### Model 2

To model Hypothesis 2, which proposes audiovisual integration at the global level, this model incorporates an integration layer at Level 2 (shown as a horizontal orange bar). Within the integration layer, the auditory and visual streams are “linked”, allowing for crossmodal sequences to be considered for SP.

#### Model 3

To model Hypothesis 3, which proposes audiovisual integration at both the local and global levels, this model incorporates integration layers at both Levels 1 and 2. The integration layer at Level 1 allows for crossmodal A-to-V and V-to-A transitions to be considered in TP.

### Theoretical Values of Prediction and Prediction-Error Signals

Next, we examined the steady-state values of that each signal would reach once TP and SP have been learned. Importantly, our analysis focused only on the signals generated by the last stimulus in each sequence, since all sequences in a block shared the same first two stimuli and crossmodal transitions occurred only at the last stimulus. We assessed the steady-state values of the prediction signals (P1_A_, P2_A_, P1_V_, and P2_V_) for the last stimulus with the principle outlined by Chao et al. (2022): The optimal value for each prediction signal is determined by minimizing the mean squared prediction error within the corresponding neural population. Utilizing this principle, once determining TP and SP, we can derive the values for all relevant prediction and prediction-error signals for each sequence (see further details in Methods).

To further elucidate model values in three models, we use sequences AAV and AAA in Block 1 as illustrative examples (Figure 2C) (comprehensive model values for all blocks can be found in Supplementary Figure 2). Here, we highlight several key points:

- The values of the prediction signals are identical for sequences AAV and AAA. This is because the prediction of the upcoming last stimulus does not depend on the identity of the last stimulus.
- In Models 1 and 2, local audiovisual integration is absent, and TP_A**→**V_ = 0. Therefore, the last stimulus V in AAV is unpredictable based on stimulus transitions, and local prediction P1_V_ = 0.
- In Model 1, global audiovisual integration is absent, and SP_AAV_ = 0. Therefore, the last stimulus V in AAV is unpredictable based on sequence structure, leading to global prediction P2_V_ = 0.
- In the integration layer where two streams are linked, different prediction signals are sent down in the auditory and visual streams to minimize their respective prediction errors. For example, in Model 2, the values of P2_A_ and P2_V_ differ since they aim to minimize different prediction errors (i.e., PE2_A_ and PE2_V_, respectively).

To extract the prediction error, we conducted within-block contrasts: contrasting the model values between sequences AAV and AAA (AAV – AAA) in Blocks 1 and 2, and the model values between sequences VVA and VVV (VVA – VVV) in Blocks 3 and 4. These contrasts eliminated the prediction signals, which are identical for both sequences within a block, and contained only the contrast values of sensory and prediction-error signals. An illustrative example of these contrast values in Block 1 is shown in Figure 2D.

For the integration layers, we propose that the neuronal populations from both streams would spatially cluster in the brain to facilitate effective information integration. This clustering would resemble the specialized subareas for processing specific modalities in the multisensory superior temporal sulcus (Barraclough et al., 2005; Schroeder & Foxe, 2002). In the model, neuronal populations are grouped across both auditory and visual streams to represent this clustering (Figure 2D). This is especially crucial in Model 3. In Model 3, PE2_A_ and PE2_V_ are collectively transmitted from grouped neuronal populations at Level 1 to grouped neuronal populations at Level 2. Consequently, we combined them into a single composite signal, PE2_A/V_ (= PE2_A_ + PE2_V_), based on the assumption that they cannot be spatially separated due to the limited resolution of EEG.

The comprehensive contrast values for all blocks are shown in Figure 2E. Additionally, we constructed alternative models featuring modified elements related to integrated neuronal populations and signals (see details in Supplementary Figure 1). These alternative models were subsequently evaluated.

### Model Evaluation with EEG Data

Next, we tested the three hypotheses by quantifying how well the modeled contrast values (as in Figure 2E) fit the EEG data. The EEG data were preprocessed to reduce artifacts and impact of volume conduction (see details in Methods). For each channel and trial, the event-related spectral perturbation (ERSP) was calculated (baseline: –0.2 to 0 sec, relative to the last stimulus). In trials where stimulation was applied to the left side, the channel assignments for ERSP were mirrored from left to right. This mirroring aimed to align the sensory responses with trials where stimulation was delivered to the right side and ensured the left hemisphere to serve as the contralateral side in both scenarios. After the mirroring, the ERSPs were then averaged across trials for each trial type in each block, and for each participant and each channel.

To capture prediction-error signals in accordance with the model shown in Figure 2D, we contrasted the averaged ERSPs from different trial types: AAV – AAA in Blocks 1 and 2, and VVA – VVV in Blocks 3 and 4. Significant contrast responses were identified as those significantly different from zero using bootstrap analysis (n = 30 participants, 1000 resamples, alpha = 0.05, two-tailed, with false discovery rate correction), and non-significant responses were replaced with zeros. An example of a significant contrast response is shown in Figure 3A. Figure 3B shows the overall significant contrast responses, averaged across all channels for each block. These responses were mainly observed after the last stimulus, exhibiting varying patterns across blocks (see also Supplementary Analysis 1 for further details). We then pooled all the significant contrast responses into a tensor with 3 dimensions: *Block* (4 blocks), *Channel* (60 channels), and *Time-Frequency* (450 time points × 100 frequency bins).

**Figure 3.**
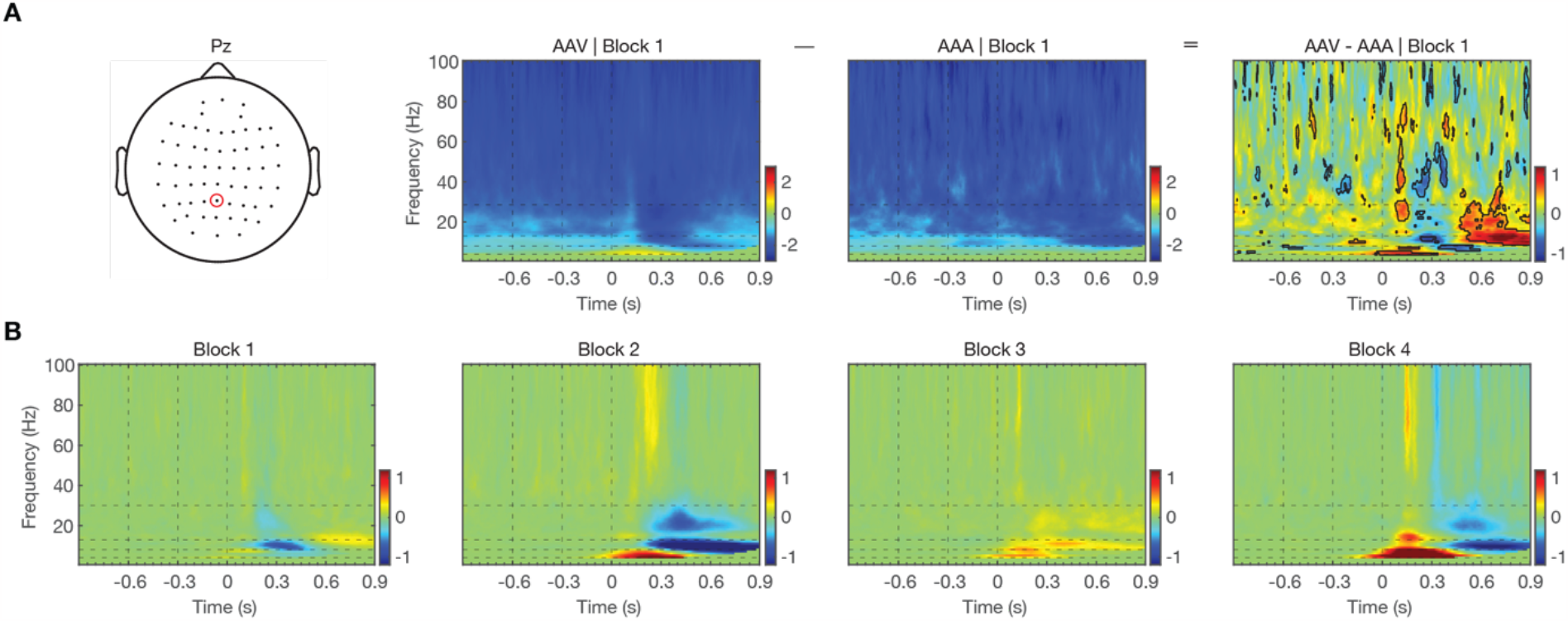
Significant contrast responses. **(A)** Example responses of AAV and AAA in Block 1 at channel Pz, and their contrast. Time zero represents the onset of the last stimulus of a sequence. The black vertical dashed lines represent the onsets of the stimuli of a sequence. The black horizontal dashed lines delineate the ranges of theta (4-8 Hz), alpha (8-13 Hz), beta (13-30 Hz) and gamma (30-100 Hz) bands. The black contours represent significant differences (bootstrap, alpha level of 0.05). **(B)** The significant contrast responses averaged across all channels in the four blocks.

To fit the 3D tensor with the models, we used a decomposition approach to break down the tensor into various components of sensory and prediction-error signals, based on their corresponding contrast values in the models. For example, in the case of Model 1, we restricted the tensor to break down into 6 designated components: S_A_, S_V_, PE1_A_, PE1_V_, PE2_A_, and PE2_V_. Each of these components was activated across all four blocks, precisely matching the activation profiles shown in Figure 2E. This was achieved by fixing the *Block* dimension to the modeled contrast values in the parallel factor analysis (PARAFAC), a high-dimensional decomposition method (Bro & Kiers, 2003; Harshman & Lundy, 1994). We performed PARAFAC 50 times, each with a random initialization.

Following this, we assessed which model offered a better explanation of the data. First, we used the core consistency diagnostic (CORCONDIA) to quantify the interaction between components (Harshman & Lundy, 1994). A higher CORCONDIA indicates less interaction between the components, indicating a more appropriate fit for the decomposition (Bro & Kiers, 2003; Pouryazdian et al., 2016). In Figure 4A, CORCONDIA was found to be 0 ± 0.1% in Model 1, 14.8 ± 8.8% in Model 2, and 83.3 ± 0.6% in Model 3 (mean ± standard deviation, n = 50 random initializations). Model 3 showed a significantly higher CORCONDIA value compared to Model 1 (t[98] = 928.55, *p*<0.001) and Model 2 (t[98] = 35.42, *p*<0.001). Additionally, we assessed the model’s goodness of fit using the Bayesian Information Criterion (BIC). A lower BIC value suggests a better balance between the model’s fit and its complexity. Model 3 showed the lowest BIC value compared to Model 1 (t[98] = -3.04e+07, *p*<0.001) and Model 2 (t[98] = - 7.62e+06, *p*<0.001). Furthermore, Model 3 outperformed the other alternative models outlined in Supplementary Figure 1, displaying both higher CORCONDIA and lower BIC values (see Supplementary Table 1).

**Figure 4.**
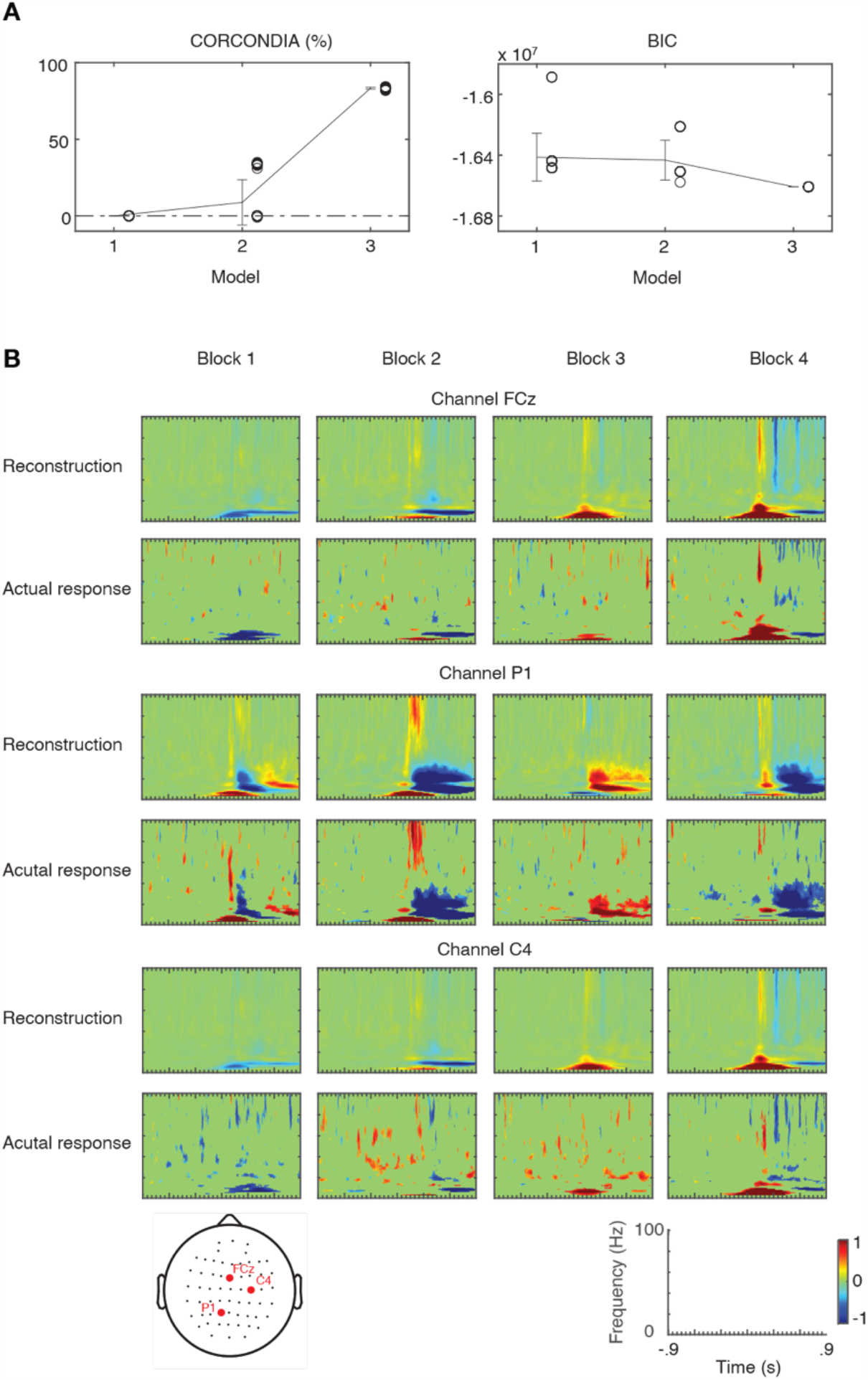
Model comparison and reconstruction. **(A)** The core consistency diagnostic (CORCONDIA) and Bayesian information criterion (BIC) gained with data fitting 50 times respectively in each of the three models. The mean and standard deviation are shown with a black error bar, and fits are shown with black circles. **(B)** The time-frequency representation in four blocks at channel FCz, P1, and C4 are reconstructed from Model 3. The actual responses are the significant contrast responses, as the comparison to the reconstructed representations.

These results consistently indicated that Model 3 was the best-fitting model. To further substantiate this, we reconstructed the significant contrast responses using the 5 components derived from Model 3 (see details in Methods) and demonstrated that Model 3 was effective in capturing the diverse responses across different channels and blocks (Figure 4B).

### Hierarchical Prediction-Error Signals Extracted From Model 3

Figure 5A shows the 5 components in Model 3 (S_A_, S_V_, PE1_A_, PE1_V_, and PE2_A/V_) extracted from PARAFAC. Each component extracted from PARAFAC is represented by the score or loading matrix (activation values) along the original dimensions: *Block, Channel*, and *Time-Frequency*. The first dimension was anchored to the 4 modeled contrast values. The remaining two dimensions captured how the component was activated across 60 channels (*Channel*), and across 450 time points and 100 frequency bins (*Time-Frequency*), describing its spatio-spectro-temporal signature.

**Figure 5.**
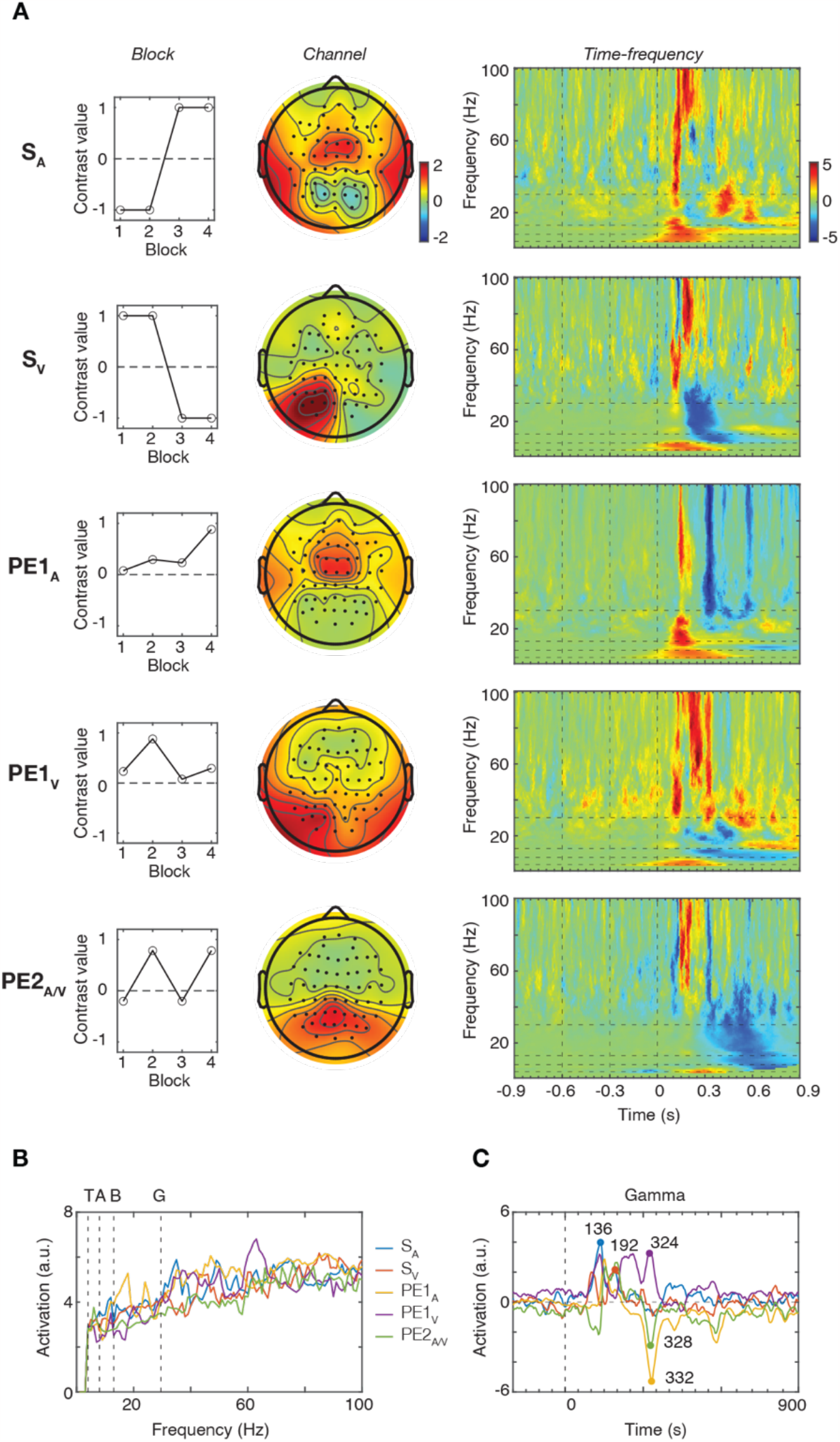
Neural components extracted from the best-fitting model. **(A)** The sensory components (S_A_ and S_V_) and the prediction-error components (PE1_A_, PE1_V_ and PE2_A/V_), showing the model contrast values across four blocks (left), the topography (middle), and the time-frequency representation (right). The time-frequency representations are normalized to equalize the strengths for visualization only by dividing the responses with the standard deviation of the corresponding frequency band. **(B)** The frequency profile of the five components. The solid vertical lines represent the range of the frequency bands, labeled on top (T: Theta; A: Alpha; B: Beta; G: Gamma). **(C)** The temporal profile of the gamma band. The peak latencies of the extracted components are shown.

S_A_, the sensory signal in the auditory stream, was found in the bilateral temporal and central areas with increases in the theta (4-8 Hz), alpha (8-13 Hz), beta (13-30 Hz), and gamma (30-100 Hz) bands after the last stimulus. S_V_, the sensory signal in the visual stream, was found in the contralateral parieto-occipital area with increases in the theta and gamma bands after the last stimulus and then decreases in the alpha, beta, and gamma bands.

PE1_A_, the first-level prediction-error signal in the auditory stream, was found in the bilateral temporal and central areas with increases in the theta, alpha, beta, and gamma bands and then a decrease in the gamma band. PE1_V_, the first-level prediction-error signal in the visual stream was found in the bilaterally parieto-occipital area with increases in the theta band and gamma bands and then decreases in the alpha and beta bands. PE2_A/V_, the second-level prediction-error signal in the integrated stream, was found in the central-parietal area with an increase in the gamma band and then decreases in the theta, alpha, and gamma bands, and a long-lasting decrease in beta band oscillations.

Spatially, S_V_ was activated more in the contralateral area due to the lateral stimulation delivery. On the other hand, S_A_ was activated in both hemispheres, which could result from less focal sound delivery by the speakers compared to headphones. For the prediction errors, PE1_V_ appeared in the bilateral scalp areas and PE1_A_ appeared in the central area, suggesting that the sensory signals located in the contralateral sensory areas now propagated to the central area and other brain areas. Spectrally, we measured the peak value for each component in each frequency band the 0-0.9 sec time window (Figure 5B). All five components exhibited higher values in the gamma band. Temporally, we focused on the dynamics of gamma-band signals by averaging the values across frequency bins within the gamma band for each component (Figure 5C). The peak latencies for S_A_, S_V_, PE1_A_, PE1_V_, and PE2_A/V_ were 136 msec, 192 msec, 332 msec, 324 msec, and 328 msec, respectively. These latencies suggested a sequential bottom-up signal propagation from sensory input to prediction error, in line with signal flows in our model. Peak latencies for other frequency bands are provided in Supplementary Figure 3.

### Crossmodal Integration During Learning

The PARAFAC results suggested that the unimodal prediction-error signals, PE1_A_ and PE1_V_, were integrated in the central-parietal area to generate the crossmodal prediction-error signal PE2_A/V_. To explore how this integration established during learning, we assessed the functional connectivity between the relevant areas. For each component (PE1_A_, PE1_V_, or PE2_A/V_), we selected the five channels exhibiting the highest values based on the PARAFAC results in the *Channel* dimension. This resulted in 25 connections between PE1_A_ and PE2_A/V_. Meanwhile, there were only 24 connections between PE1_V_ and PE2_A/V_, since there was one channel shared between them (Figure 6A). To assess learning, we divided the trials within a single block into 11 continuous time windows. Trials from each of these windows were then grouped across all four blocks.

**Figure 6.**
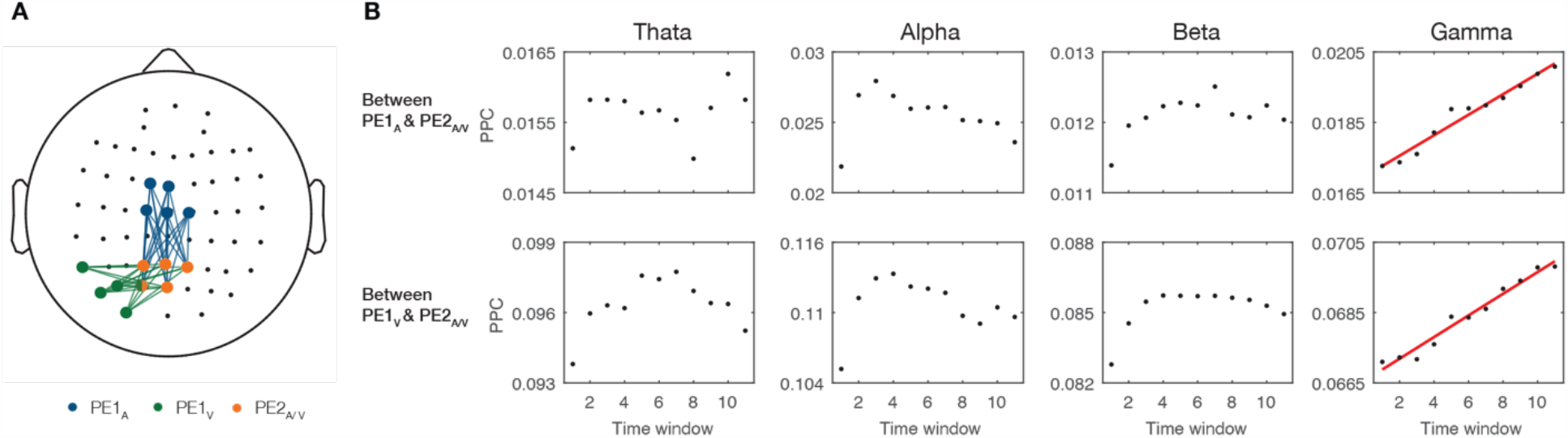
Functional connectivity during learning. **(A)** The channels selected from the prediction-error components: FC1, FCz, C1, C2 and Cz for PE1_A_ (blue dots), P7, PO3, PO5, PO7 and O1 for PE1_V_ (green dots), and P1, P2, Pz, PO3 and POz for PE2_A/V_ (orange dots). The 25 connections are paired between the channels from PE1_A_ and PE2_A/V_ (blue solid lines). The 24 connections are paired between between the channels from PE1_V_ and PE2_A/V_ (green solid lines). **(B)** For each frequency band and stream, the mean pairwise phase consistency for each window is shown with a black dot, and a significant linear trend over 11 windows (permutation, 0.05) is indicated with a red solid line.

To measure spectral functional connectivity, we used pairwise phase consistency (PPC), a method measuring phase synchronization between two signals across trials while minimizing bias related to the number of trials (Vinck et al., 2010). For each connection, time window, and participant, PPC values were calculated and then averaged across frequency bins within 4 distinct frequency bands (theta, alpha, beta, and gamma). These averaged PPC values were further averaged across the relevant connections (either 25 or 24 depending on the components involved), across time points (0-0.9 sec after the last stimulus), and across participants. For the connections between PE1_A_ and PE2_A/V_, 11 PPC values were generated for each frequency band to illustrate how functional connectivity evolved over the course of the trials (learning stages) (Figure 6B). Similarly, between PE1_V_ and PE2_A/V_, 11 PPC values were obtained for each frequency band to depict changes of functional connectivity.

Through linear regression analysis, a significantly positive slope was found in the gamma band between PE1_A_ and PE2_A/V_ (*p* = 0.001, permutation, n = 1000 shuffles, alpha = 0.05, two-tailed), and between PE1_V_ and PE2_A/V_ (*p* = 0.007). These results indicated that the gamma-band interactions between the central-parietal area and the sensory areas gradually strengthened during learning and suggested that the central-parietal area gradually emerged as a hub to integrate crossmodal information.

## Discussion

We combined a crossmodal experimental paradigm with a hypothesis-driven analysis to examine hierarchical crossmodal predictive coding for sequence processing. We demonstrated that hierarchical crossmodal predictions can be established by learning TP and SP in a higher-order brain region that integrates information from different sensory streams. Our finding provides a generalized framework for crossmodal predictive coding and reveals neural signatures of its representation and learning.

### Error-driven convergence

From the predictive coding perspective, convergence of sensory inputs along the hierarchy plays a critical role to minimize prediction errors. To demonstrate this, we calculated the theoretical values of the overall prediction error for each block in each model; that is the minimum of mean-squared prediction error, the principle in model value calculation. For each block (e.g., Block 1), the squared values of PE1_A_, PE1_V_, PE2_A_, and PE2_V_ of each trial type (e.g., AAV or AAA) were weighted by the probability of its occurrence and then summed together. This summed value represents the overall prediction error, which was the same between Blocks 1 and 3, since the model values were the same in these two blocks with just the two sensory streams swapped. Similarly, the overall prediction error was the same between Blocks 2 and 4. The overall prediction errors in Model 3 were found to be the smallest among the three models (Supplementary Figure 4). Convergence early at functional level of stimulus-to-stimulus transitions enables the brain to achieve the smallest sum of prediction errors, which is in line with the general framework of predictive coding theory.

### A third area for convergence

The central-parietal area neither located in the auditory central area nor visual occipital area may has access to merge crossmodal information, suggested by the results of functional connectivity and frequency oscillations. The connectivity strength between the central-parietal area and the two sensory areas increased during learning, particularly in the gamma frequency, which is thought to mediate bottom-up transmissions of prediction errors from a lower neuronal layer to a higher neuronal layer (Bastos et al., 2015; Fries, 2005; Michalareas et al., 2016). Also, the temporal order of the peak gamma oscillations evoked in two primary sensory areas (S_A_ and S_V_) and then in the central-parietal area (PE2_A/V_) further provides directional support where information may be carried from sensory to higher areas over time. In addition to the gamma oscillation, the component of PE2_A/V_ contained a late long-lasting negative beta-band component, which shares similarity with reduced beta-band power found in previous research using multimodal tasks (Göschl et al., 2015; Roa Romero et al., 2015). Also, the desynchronization of the beta oscillation suggests a top-down transmission to update predictions for further error minimization (Arnal & Giraud, 2012; Chao et al., 2018; Jiang et al., 2022).

Those findings imply that this third area receives audiovisual feedforward information carried via the gamma oscillations, and such information would be used to reset the state for precise expectations of incoming information via the desynchronization of the beta oscillations.

However, we also observed theta and alpha oscillations evoked around the onset of the gamma oscillation in the components of two sensory inputs and first-level prediction errors, while we observed reduced theta and alpha oscillations and thus less coupling between the low and high frequency oscillations in the components of the second-level prediction error. Evidence of phase-amplitude cross-frequency coupling has been discovered to connect different brain regions and have large-scale networking between regions such as the hippocampus and prefrontal areas (Bragin et al., 1995; Canolty et al., 2006; Cohen et al., 2009; Sirota et al., 2008). Further advanced recording and methodologies such as Electrocorticography (ECoG) in the hippocampus area and whole cortical surface can be combined with cross-frequency coupling analysis to verify if lower- and higher-hierarchy crossmodal processing involves different ranges of distributed networks.

Although the central-partial area plays a plausible role, studies have shown that convergence of sensory processing across modalities can occur already in subcortical and lower cortical areas. For example, inferior and superior colliculus have both been found to receive inputs of different sensory modalities (Champoux et al., 2006; Meredith & Stein, 1983). This could also be achieved by interconnections between sensory-specific primary cortices, which have been identified between the auditory and visual areas, and between auditory and somatosensory areas (Driver & Noesselt, 2008; Schroeder et al., 2001; Schroeder & Foxe, 2002). We suggest that early convergence could occur when crossmodal transitions are processed at a short timescale, such as visual speech occurring a few hundred milliseconds ahead and leading to the prediction of acoustical input of speech spectrograms and phonetic features (O’Sullivan et al., 2021). When considering complex sequences with multiple time scales, probability computations would be operated at a more distributed brain network. Our study used arbitrary stimuli to create two-level temporal regularities, while real-world stimuli can consist of more than two timescales of regularities, e.g. in music and language (Dehaene et al., 2015). It is needed to further reveal this third area or a larger brain network processing complex sequences with natural stimuli and thus generalize crossmodal predictive coding in the multi-level hierarchy.

### Limitations and future work

There is still room for further investigation of the hierarchical crossmodal predictive coding model, including neural representations of crossmodal prediction signals, trial-by-trial prediction updates, and interpretation of atypical audiovisual integration. Firstly, crossmodal prediction signals were not estimated directly. We modeled the prediction-error signals when established crossmodal predictions were violated, and degrees of prediction errors also reflected crossmodal predictability. To extract neural representations of prediction signals in order to construct a comprehensive predictive coding network, we would need to break the interdependence between crossmodal predictions and crossmodal prediction errors by considering their timing delay. That is, a prediction is activated before input is received and then a prediction error is evoked after input is received.

Secondly, while we could provide some estimate of the learning effect by dividing EEG trials in one block into several segments, our model is limited by design to steady-state values, and trial-by-trial dynamics of crossmodal prediction updates over time remains unknown. Although Hierarchical Gaussian Filter is a way to measure the dynamics of how prediction errors are optimized at different levels (Iglesias et al., 2013; Mathys et al., 2014), the hierarchical structure cannot be shaped when different timescales of statistical regularities are learned and affect hierarchical interactions. Still, our mode with the hierarchical structure and analyses could be greatly improved with the explicit inclusion of learning over time.

Finally, altered multisensory integration has been used to explain the difficulties of involving in audiovisual communication experienced by individuals with autism spectrum disorder (Meyer & Noppeney, 2011; Saalasti et al., 2011). In contrast, multisensory integration can allow older people to compensate for declines in physical functions (e.g., reduced visual or auditory acuity) by inferring information from other sensory sources (Freiherr et al., 2013). Our task with a model approach can provide a platform to evaluate multisensory integration from a hierarchical view and offer insights into the neural mechanism about how and why deficits or gains in multisensory integration happen.

## Materials and Methods

### Participants

Thirty participants were recruited for this study (15 males and 15 females; age = 22.0 ± 2.4 years). The inclusion criteria were: (1) age between 20-50 years, (2) no difficulty in understanding the experimental procedure due to apparent deficits in cognition, hearing and vision, and (3) no medical history of neurological or psychological diagnosis. Signed consent was received from the participants before the experiment, and all protocols were approved by the research ethics committees of The University of Tokyo (No. 21-372) and National Taiwan University Hospital (No. 201906081RINA).

### Stimuli and task

The auditory stimulus (A) was created with three superimposed sinusoidal waves (500, 1000 and 1500 Hz; volume = 55 dB), and the visual stimulus (V) was a 16*16 green-black checkerboard (size: 4 degree visual angle with a 70 cm viewing distance) presented against a gray background (1920*1220 pixels). These two stimuli were arranged to create 4 sequences: AAA, AAV, VVV, and VVA with 200-ms duration and 300-ms SOA. A random interval of 1600-1900 ms (50ms as a step) was inserted between the offset of one sequence and the onset of the next sequence

Four blocks were respectively assigned a frequent sequence and an infrequent sequence, and constructed as follows: The 120 trials were notionally divided into four phases of 30 trials each. Each phase was built from 25 frequent and 5 infrequent sequences in pseudorandom order, yielding an even distribution of trial types within the block. Each block was presented twice, once for the left stimulation delivery, where the visual stimuli were delivered 5.28 degree visual angle away from the screen center and the auditory stimuli were delivered from the left speaker, and once for the right stimulation delivery. The order of the total eight runs was random, with a break between runs. A white fixation dot was presented at a visual angle of 3.85 degrees below the screen center. The experimental protocol was programmed with the MATLAB-based PsychtoolBox (Kleiner et al., 2007).

### Probability calculation

The values of TP and SP differ across Blocks and Hypotheses. Here, we first demonstrate the calculations of TP and SP for Block 1 in Hypothesis 1. For a single sequence AAA, there are two A-to-A transitions, and one A-to-O transition. For a single sequence AAV, there are one A-to-A transitions, and one A-to-O transition. Therefore, the total number of A-to-A transitions (denoted as TN_A**→**A_) is 2*20 (20 trials of sequence AAA) + 1*100 (100 trials of sequence AAV) = 140.

Similarly, the total number of A-to-O transitions (denoted as TN_A**→**O_) is 1*20 (sequence endings for 20 trials of sequence AAA) + 1*100 (100 trials of sequence AAV) = 120. Since there is no A-to-V transition in Hypothesis 1, the total number of A-to-V transitions (denoted as TN_A**→**V_) is zero.

TP_A**→**A_, TP_A**→**V_, and TP_A**→**O_ then can be calculated by dividing the number of corresponding transitions by the total number of transitions (= 140 + 120 + 0 = 260), which results in 140/260, 0, and 120/260, respectively (as shown in Table 1). For SP, there are 20 trials of AAA, 0 trial of AAV (no global integration), and 100 trials of AA. Thus, SP_AAA_, SP_AAV_, SP_AA_ are 20/120, 0, 100/120, respectively. The comprehensive calculations are formulized below.

For Block 1 in Hypothesis 1:

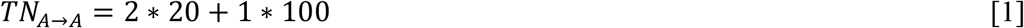

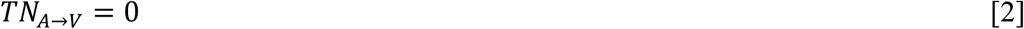

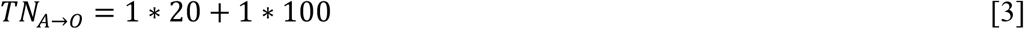

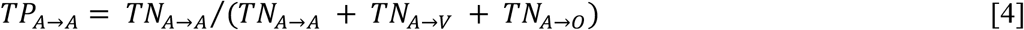

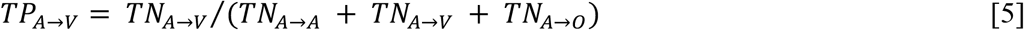

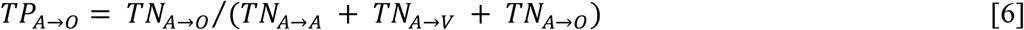

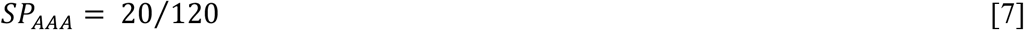

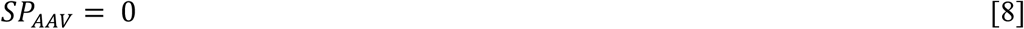

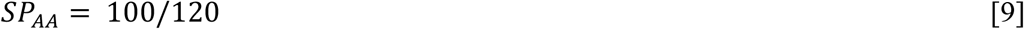

In Hypothesis 2, TPs are calculated based on Equations [1∼6] as in Hypothesis 1. For SP, since there is global integration for sequence AAV, SP_AAA_, SP_AAV_, SP_AA_ are 20/120, 100/120, and 0 respectively. In Hypothesis 3, SP are calculated as in Hypothesis 2, and TPs are calculated as in Hypothesis 1, but with different transition numbers. With local integration, TN_A**→**A_ is still 140, but TN_A**→**V_ is 1*100 (100 trials of sequence AAV) = 100, and TN_A**→**O_ is 1*20 (sequence endings for 20 trials of sequence AAA) = 20.

For Block 2, the only difference from Block 1 is the number of trials. In this block, there are 100 trials of AAA and 20 trials of AAV. The calculations of TP and SP can be conducted according to Equations [1∼15], with the numbers 100 and 20 being swapped.

For Blocks 3 and 4, the only difference from Blocks 1 and 2 is the modality. The calculations of TP and SP can be conducted based on the same principle, but with the modality A and V being swapped. For example, TP_A**→**V_ is swapped to TP_V→A_, and SP_AAV_ is swapped to SP_VVA_.

### Model value calculation

The model values were calculated from TP and SP based on Chao et al., 2022. To demonstrate this, we use Block 1 as an example (see Figure 2C). For the A-to-A transition (with the probability of TP_A→A_), S_A_=1, and thus PE1_A_ = |1 – P1_A_| (|.| represents the absolute value). For the other transitions (A-to-O or A-to-V, the probability is 1 – TP_A→A_), S_A_=0, and thus PE1_A_ = |0 – P1_A_|. Therefore, the corresponding mean-squared error, denoted as MSE1_A_, is:

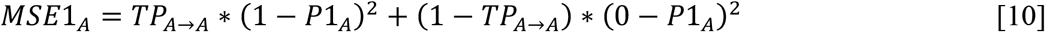

which is a function of P1_A_ and can be minimized by P1_A_ = TP_A→A_. A similar calculation can be applied to Level 2, where PE2_A_ = |PE1_A_ – P2_A_|. For sequence AAA (with the probability of SP_AAA_), S_A_=1 and PE1_A_ = |1 – P1_A_|, and thus PE2_A_ = ||1 – P1_A_|– P2_A_|. For the other sequences (AA or AAV, the probability is 1 – SP_AAA_), S_A_ = 0 and PE1_A_ = |0 – P1_A_|, and thus PE2_A_ = ||0 – P1_A_|– P2_A_|. Therefore, the corresponding mean-squared error, denoted as MSE2_A_, is:

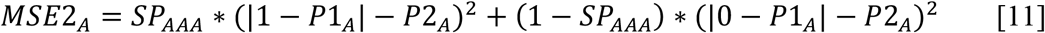

which is a function of P2_A_ and can be minimized by P2_A_= SP_AAA_*(|1 – P1_A_|) + (1– SP_AAA_)*P1_A_. For P1_V_ and P2_V_, similar calculations can be applied with the replacement from TP_A→A_ to TP_A→V_ and from SP_AAA_ to SP_AAV_. Furthermore, the same calculations can be applied to Block 2 but with different values of TP and SP. For Blocks 3 and 4, similar calculations are done by replacing TP_A→A_ with TP_V→A_, TP_A→V_ with TP_V**→**V_, SP_AAA_ with SP_VVA_, and SP_AAV_ with SP_VVV_. The comprehensive calculations are shown below and provided in GitHub (<u>link</u>). For different hypotheses, different TP and SP values are used.

For Blocks 1 and 2:

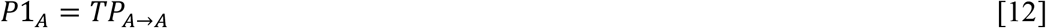

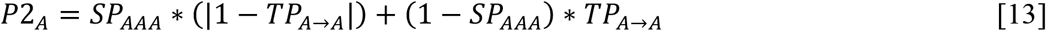

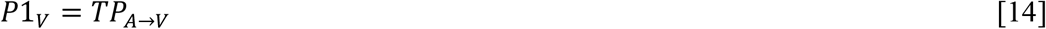

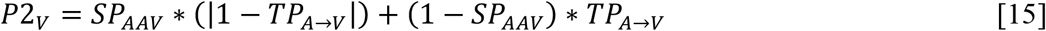

For Blocks 3 and 4:

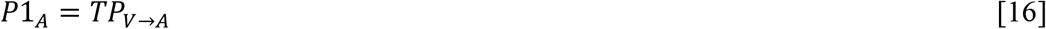

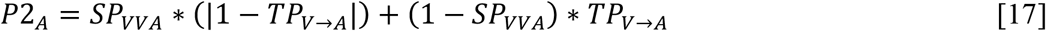

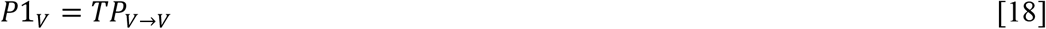

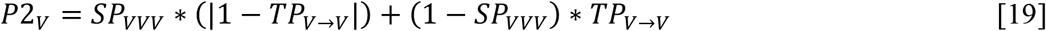

Once, the predictions are determined and the prediction errors are calculated as:

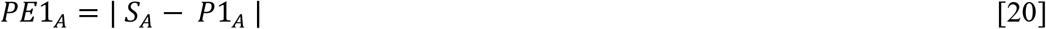

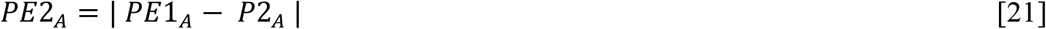

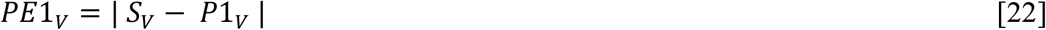

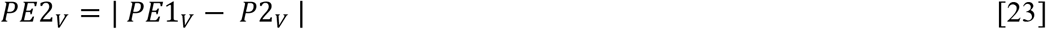

### EEG acquisition and preprocessing

EEG was recorded by a 64-channel 10-20 system Quick Cap and Neuroscan SynAmps RT amplifier (Compumedics Neuroscan USA, Ltd. Charlotte, NC, USA). The impedances were kept below 10 kΩ for the two eye electrodes and below 5 kΩ for the rest of the electrodes. The signals were collected with an online reference electrode near Cz, a 500 Hz sampling rate, and an online band-pass filter between 0.01 to 100 Hz. Event codes were sent at the onset of the first stimulus in a sequence. We used EEGLAB (Swartz Center for Computational Neuroscience) (Delorme & Makeig, 2004) for data preprocessing including extracting epochs from -1 to +1.9 sec relative to the onset of the first stimuli (function: pop_epoch.m), and manually removing bad epochs when there was an obvious movement artifact or 80% of amplitudes were over ± 100 microvolts.

Then, components containing muscular and eye artifacts were removed using independent component analysis (ICA) (function: pop_ica.m) with ADJUST toolbox (Mognon et al., 2011) (function: interface_ADJ.m). Current source density (CSD) analysis was conducted using CSD Toolbox to reduce the impact of volume conduction (function: CSD.m) (Kayser & Tenke, 2006). For CSD calculation, the montage of spherical coordinates was created using the channel information of the Quick Cap without channels CB1 and CB2 (function: Get_GH.m). The smoothing constant lambda was set as 1e-5, and the head radius was set as 10 cm.

### Event-related spectral perturbation (ERSP) and significant contrast responses

We used Fieldtrip Toolbox (Oostenveld et al., 2010) to quantify the time-frequency representation (from 1 to 100 Hz) of the preprocessed EEG for each trial, channel, and participant. The Morlet wavelet transformation was used with the wavelet cycles of 7 (function: ft_freqanalysis.m). To measure the event-related spectral perturbation for each trial, we performed decibel baseline normalization with the baseline period from –0.2 to 0s (time zero as the onset of the first stimulus) (function: ft_freqbaseline.m).

To make the left hemisphere the contralateral side for both left-side and right-side stimulation delivery, for trials where stimulation was applied to the left side, the ERSPs’ channel assignment was mirrored from left to right, and for trials where stimulation was applied to the right side, the channel layout was unchanged. Then, the ERSPs were then averaged across trials for each trial type in each block, and for each participant and each channel.

For each participant and each channel, we contrasted the averaged ERSP in four blocks. The significance of the contrast across subjects was estimated using bootstrap analysis (function: bootci.m).

### PARAFAC analysis

PARAFAC analysis is a generalization of the principal component analysis to higher-order arrays. This method decomposes a tensor into a sum of outer products of vectors, which helps in identifying underlying patterns or structures in the multi-dimensional data. We performed the analysis using the N-way toolbox (Andersson & Bro, 2000) (function: parafac.m). The input was the significant contrast responses (4 *Block* × 60 *Channel* × 450*100 *Time-Frequency*). The option of “Fixmode” was applied to *Block* with the modeled contrast values while no constraint was applied on dimensions of *Channel* and *Time-Frequency*. The convergence criterion was set to be 1e-6. We repeated the decomposition 50 times, each time with a set of random orthogonalized values for initialization. Each component decomposed from PARAFAC can be described by score or loading matrices (activation values) in the original dimensions: 4 model contrast values in the *Block* dimension, 60 activation values in the *Channel* dimension, and 450*100 activation values in the *Time-Frequency* dimension.

To evaluate the decomposition a CORCONDIA of 80∼90% is considered a good consistency, and values below 50% indicate a problematic model with high interaction between components (Bro & Kiers, 2003; Pouryazdian et al., 2016). On the other hand, BIC was calculated to correct the residual sum of squares (RSS) with the complexity of the model:

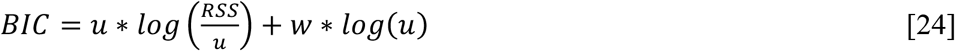

Where *u* represents the size of inputs (e.g., 4 *Block* × 60 *Channel* × 450*100 *Time-Frequency*) and *w* represents the size of estimated outputs. See the values of *w* for the three models in Supplementary Table 1.

From 50 decompositions (each with random initialization), we acquired 50 CORCONDIA values and 50 BIC values. These values were compared between models using two-sample t test, and alpha was set as 0.05 with Bonferroni correction (function: ttest2.m).

### Response reconstruction

We reconstructed the responses from 5 extracted components (i.e., S_A_, S_V_, PE1_A_, PE1_V_, and PE2_A/V_) with dimensions of *Block, Channel*, and *Time-Frequency*. The response for Block *i* and Channel *j* was reconstructed (denoted as *Recon*) as follow:

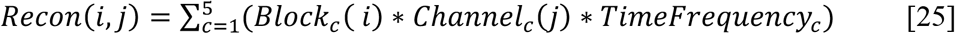

Where *c* represents the component index, where c =1 (S_A_), 2 (S_V_), 3 (PE1_A_), 4 (PE1_V_), and 5 (PE2_A/V_). For component *c, Block*_*c*_ represents the 4 model contrast values, *Channel*_*c*_ represents the 60 activation values in the *Channel* dimension, and *TimeFrequency*_*c*_ represents the 450*100 activation values in the *Time-Frequency* dimension.

### Connectivity analysis

For the spectral functional connectivity, we performed PPC with Fieldtrip Toolbox (function: ft_connectivityanalysis.m). We paired 25 connections between PE1_A_ and PE2_A/V_, and 24 connections between PE1_V_ and PE2_A/V_ because of one shared channel. We then segmented EEG responses in each block into 11 learning stages using a moving average approach. N is the number of trials in a phase of a block after bad epochs were excluded, and we created three segments (N/3 trials) for one phase and thus twelve segments in total for one block. Then, we used a window width of two segments and a step size of one segment to have 11 learning stages. For example, in a clean block (no bad trials, N = 30 from 120/4), and a segment has 10 trials (from 30/3). The first stage would include trials from 1^st^ to 20^th^ (first and second segments), and the second stage would include trials from 11^th^ to 30^th^ (second and third segments). For each stage, the trials from the four blocks were then pooled together.

PPC was evaluated at 450 time points and 100 frequency bins (1∼100 Hz). These PPC values were then averaged across frequency bins for 4 frequency bands (theta: 4-8 Hz, alpha: 8-13 Hz, beta: 13-30 Hz, and gamma: 30-100 Hz), across time points (0-0.9 sec relative to the last stimulus), across connections (25 or 24 between two components), and across participants. After average, there were 11 PPC values of learning stages for each pair of connected components and each frequency band. The linear trend of these 11 PPC values was quantified by linear regression analysis (function: fitlm.m). To test whether the slope was significantly different from zero, we performed a permutation analysis by shuffling the temporal order of learning stages in each participant and re-calculating the slope (n = 1000 shuffles, alpha = 0.05, two-tailed).

## Supporting information

all_materials

## Data availability

The raw EEG data are available from the corresponding author upon request.

## Contributions

Z.C.C. conceptualized the study. Y.T.H., C.W., and Z.C.C. designed the experimental protocol, and Y.T.H., Y.M.F., and C.F. collected the data (Y.M.F. and C.F. contributed equally). Y.T.H. and Z.C.C. proposed the theoretical models and performed the data analyses. Y.T.H wrote the manuscript, and Z.C.C., C.W., and S.K. helped with editing. All authors contributed to and have approved the final paper.

## Acknowledgments

We thank Kai-Jie Liang and Shih-Yao Mao for helping with participant recruitment and experiment preparation, and Felix B. Kern for proofreading. We also thank Hui-Fen Mao and Shih-Yao Mao for their help in managing IRB, and Shih-Yao Mao. This work was supported by Japan Society for the Promotion of Science, Japan (to Y.T. H.), World Premier International Research Center Initiative (WPI), MEXT, Japan (to Z.C.C.), and the Ministry of Science and Technology of Taiwan (MOST 109-2410-H-002-106-MY3) (to C.W. and H.M.).

