## Supplementary material for "Crossmodal Hierarchical Predictive Coding for Audiovisual Sequences in Human Brain": all_materials

Supplementary Table 1. **Model fitness across models**

| Model name | Model streams | Neuronal populations | Number of components | CORCONDIA (mean ± sd) | RSS (mean ± sd) | *w* | BIC (mean ± sd) |
|---|---|---|---|---|---|---|---|
| Model 1 | A, V | $A_0$, $A_1$, $A_2$, $V_0$, $V_1$, $V_2$ | 6 | 0 ± 0.1 | 2.1768e+5 ±7.6559 | (60 + 450*100 + 4) *6 | -1.6404e+07 ±1.5690*e+05 |
| Model 2 | A, V | $A_0$, $A_1$, $A_2$, $V_0$, $V_1$, $V_2$ | 6 | 14.8 ± 8.8 | 2.1678e+5 ±6.1386 | (60 + 450*100 + 4) *6 | -1.6454e+07 ±1.3087*e+05 |
| Model 3 | A, V | $A_0$, $A_1$, $A_2$, $V_0$, $V_1$, $V_2$ | 5 | 83.3 ± 0.6 | 2.2333e+5 ±0.0002 | (60 + 450*100 + 4) *5 | -1.6608e+07 ±3.8178 |
| Alternative Model 2 | A, V, I | $A_0$, $A_1$, $V_0$, $V_1$, $I_2$ | 6 | 36.8 ± 16.5 | 2.1413e+5 ±1.4389 | (60 + 450*100 + 4) *6 | -1.6488e+07 ±3.1375*e+04 |
| | A, V, I | $A_0$, $A_1$, $A_2$, $AI_2$, $V_0$, $V_1$, $V_2$, $VI_2$ | 8 | -0.3 ± 0.2 | 1.9685e+5 ±140.9814 | (60 + 450*100 + 4) *8 | -1.6249e+07 ±3.3283*e+03 |
| | A, V, I | $A_0$, $A_1$, $A_2$, $V_0$, $V_1$, $V_2$, $I_2$ | 8 | 2.7 ± 3.3 | 1.9872e+5 ±1.5496*e+3 | (60 + 450*100 + 4) *8 | -1.6205e+07 ±3.6231*e+04 |
| Alternative Model 3 | A, V, I | $A_0$, $A_1$, $A_2$, $AI_1$, $AI_2$, $V_0$, $V_1$, $V_2$, $VI_1$, $VI_2$ | 9 | 0.2 ± 0.5 | 1.8993e+5 ±783.9910 | (60 + 450*100 + 4) *9 | -1.6100e+07 ±1.9199*e+04 |
| | A, V, I | $A_0$, $A_1$, $A_2$, $V_0$, $V_1$, $V_2$, $I_1$, $I_2$ | 6 | 0 ± 0.1 | 2.2131e+5 ±1.1126*e+4 | (60 + 450*100 + 4) *6 | -1.6339e+07 ±2.2544*e+05 |

* CORCONDIA: core consistency diagnostic. RSS: residual sum of squares. BIC: Bayesian information criterion. sd: standard deviation. A: auditory stream/ neuronal populations; V: visual stream/ neuronal populations; I: integration stream/ neuronal population.

Supplementary Analysis 1.

To examine whether significant contrast responses were influenced by different predictability of local and global regularities, we further compared them among the four blocks. We averaged the significant contrast responses across time points after the last stimulus and across frequency bins for 4 frequency bands: theta (4-8 Hz), alpha (8-13 Hz), beta (13-30 Hz) and gamma (30-100 Hz). For each frequency band, one-way ANOVA was applied to the averaged responses with the null hypothesis that the averaged significant responses were the same across all blocks at all channels (alpha was set as 0.05) (function: anova_test, R version 4.2.2.).

The ANOVA on the responses after the last stimulus revealed that the significant contrast responses differed significantly across blocks in all frequency bands: theta band ($F[1.17, 68.96] = 38.24$, $p<0.05$), alpha band ($F[1.36, 80.48] = 76.28$, $p<0.05$), beta band ($F[1.26, 74.34] = 42.08$, $p<0.05$), and gamma band ($F[1.97, 116.36]= 91.05$, $p<0.05$).

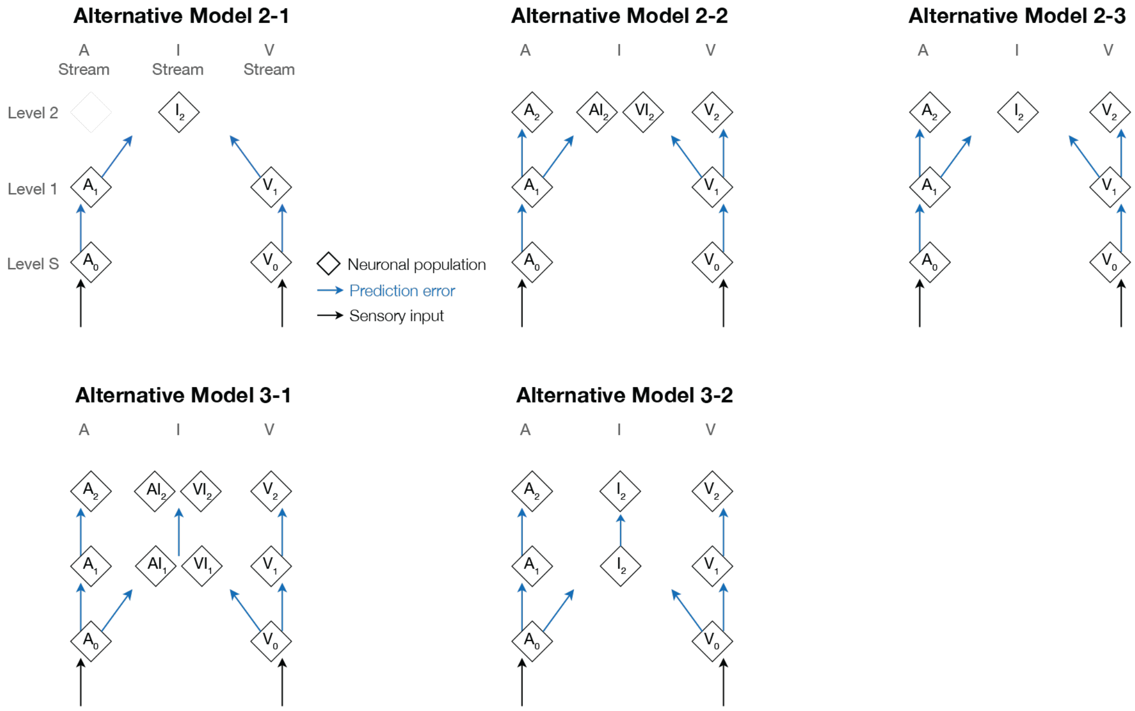


Supplementary Figure 1. **Alternative models.** Here, we show neuronal populations (white diamonds) and signal flows of prediction errors (blue solid upward arrows) and sensory inputs (black solid upward arrows) after within-block comparisons. There are two distinct differences from the three models. First, the neuronal populations, $I_1$ and $I_2$, were at the integration stream (I) and sending the same prediction to the two sensory streams (A and V) or the integration stream. Secondly, auditory and visual sensory inputs not only integrated at the I stream but also continued to transmit to their corresponding stream.

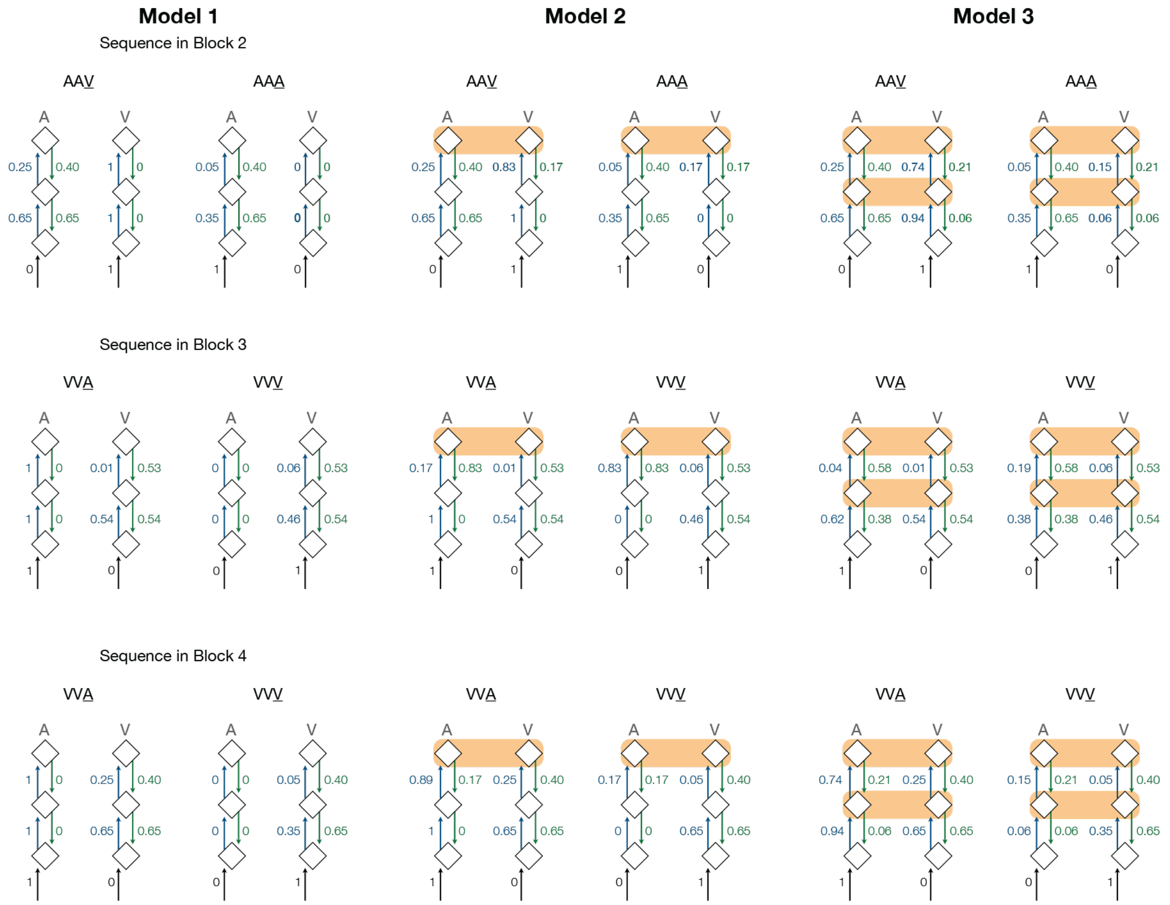


Supplementary Figure 2. **Model values in Blocks 2, 3 & 4 in Models 1, 2 & 3.** The notation of the model structures and representations is the same as Figure 2B.

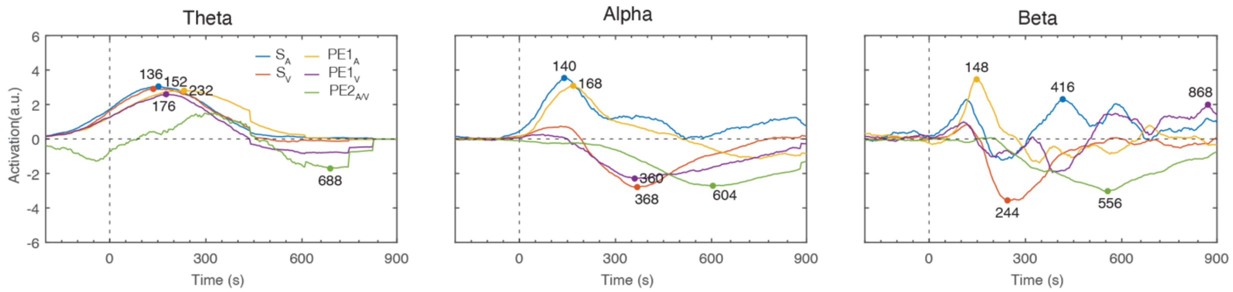


Supplementary Figure 3. **The temporal profile of theta, alpha and beta bands.** The frequency ranges of the theta, alpha, beta bands are 4-8 Hz, 8-13 Hz and 13-30 Hz respectively. The peak latencies of the extracted components are shown.

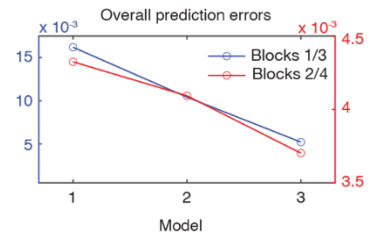


Supplementary Figure 4. **The theoretical values of the overall prediction errors.** The values for Blocks 1 and 3 are shown in blue against the y-axis on the left, and the values for Blocks 2 and 4 are shown in red against the y-axis on the right.
